# Evolution of the Spatial transcriptomic landscape during the progression of high-grade pancreatic intraductal papillary mucinous neoplasms to invasive cancer

**DOI:** 10.1101/2024.08.24.608470

**Authors:** Sirui Peng, Qiangxing Chen, Zixin Chen, Mengling Yao, Yunqiang Cai, Du He, Yu Cai, Ke Cheng, Jun Li, He Cai, Pan Gao, Xiafei Gu, Xin Wang, Yongbin Li, Man Zhang, Lingwei Meng, Qi Xia, Panpan Xu, Jin Zhou, Zhong Wu, Bing Peng

## Abstract

Intraductal papillary mucinous neoplasms (IPMN) is one of the known precancerous lesions. Patients’ prognosis is aggravated as IPMN transforms into invasive Pancreatic Ductal Adenocarcinoma (PDAC). The molecular mechanisms underlying this progression lack effective experimental models and urgently need to be elaborated. We performed spatial transcriptomics (ST) on fresh tissue samples from the same patient including normal pancreas, high-grade IPMN, and invasive PDAC, and described the step-by-step development of transcriptional landscape including clone evolution and adjacent TME feature variation. Our findings identified the master transcript factors and critical signaling pathways promoting IPMN progression to invasive PDAC. Additionally, both IPMN and PDAC harbored the ELF3, MYC, and KLF4 amplification. The Spatial CNV profile demonstrated significant heterogeneity among PDAC in their spatial distribution compared to IPMN, with seven distinct subclones showing diverse functions, such as hypoxia, oxidative phosphorylation, and epithelial-mesenchymal Transition. We observed a marked shift in the immune landscape, with the depletion of CD4+ T-cells and dendritic cells and an increase in immune-suppressive M2 macrophages in invasive PDAC, indicating a transition to an immune-evasive microenvironment. Additionally, cancer-associated fibroblasts (CAFs), particularly myofibroblastic CAFs, were enriched adjacent to invasive PDAC, suggesting their active role in tumor progression. By leveraging spatial transcriptomic analysis, our study provides comprehensive insights into the intricate molecular landscape that underlies the progression of IPMNs to invasive PDAC. These findings not only enhance our understanding of this complex process but also offer valuable knowledge for early diagnosis and intervention.

**Highlights:**

- Spatial CNV analysis reveals clonal evolution and distinct subclones in PDAC.
- Key drivers like ELF3, MYC, and KLF4 are amplified in both IPMN and invasive PDAC.
- Immune landscape shifts from pro-inflammatory in IPMN to immune-evasive in PDAC.
- Enrichment of myofibroblastic CAFs suggests their role in tumor progression

## Introduction

Intraductal papillary mucinous neoplasms (IPMN) are recognized as bona fide precursor lesions of Pancreatic adenocarcinoma (PDAC), which is projected to become the 2^nd^ leading cause of cancer-related mortality worldwide^1^. Effective treatment of PDAC hinges on a deep understanding of its molecular pathophysiology. According to the Revised Terminology (2015) for the grading of dysplasia^2^, IPMNs are classified as low-grade (LG) or high-grade (HG) dysplasia, with the latter having a greater propensity for aggressive progression, including IPMN with invasive carcinoma. IPMNs can originate from branch ducts (BD-IPMN), the main duct (MD-IPMN), or both (mixed type), and they encompass three distinct histopathological subtypes: gastric (most common, typically arising from branch ducts), intestinal, and pancreaticobiliary. The intestinal and pancreaticobiliary subtypes can emerge in either the main or branch ducts and are usually associated with high-grade IPMN. It is well-established that invasive carcinoma can arise from IPMN^3^, with 57% to 92% of patients with main-duct IPMN reporting malignant transformation in situ, compared to less than half of those with branch duct involvement^4^. Given the distinct biological behaviors and varying risks of malignant transformation, a deeper understanding of pancreatic biology and disease progression is crucial for the discovery of biomarkers and molecular targets for cancer interception.

Molecular studies have shown that the malignant transformation of IPMN is linked to the accumulation of genetic alterations. Early driver mutations in oncogenes such as KRAS and GNAS are well-documented, while mutations in tumor suppressor genes like RNF43, CDKN2A, and TP53 are typically acquired during later stages of tumorigenesis^5,6^. Recently, single-cell transcriptomic technology has been employed to unravel the pathophysiology of IPMN progression, offering unprecedented insights into tumor heterogeneity and the diverse microenvironment^7^. However, since single-cell RNA sequencing (scRNA-seq) lacks spatial context, interpreting the spatial crosstalk that could provide critical insights into tumor evolution and microenvironment interactions remains challenging. Spatial transcriptomic (ST) sequencing technology, developed by 10X Genomics, has been applied to better understand the spatial transcriptional landscape of whole tumors in various cancer types^8,9,10^.

In this study, we collected paired samples of normal pancreas, high-grade IPMN, and invasive pancreatic ductal adenocarcinoma (PDAC) from the same patient. To comprehensively elucidate the spatial evolutionary trajectory from IPMN to invasive PDAC, we performed ST on these three samples and employed multi-analytical approaches to identify critical transcription factors involved in dynamic disease progression, novel intra-tumoral clonal heterogeneity, and the impact of microenvironmental differences between IPMN and PDAC. We believe this work will provide crucial molecular insights into the mechanisms underlying PDAC evolution and contribute to discovering novel early diagnosis biomarkers and potential therapy targets.

## Materials and Methods

### Clinical Fresh Samples Collect and Process

We collected three fresh samples (C15, C16, and C17, respectively) from a female patient diagnosed with a pancreatic tumor (Figure 1A). Briefly, the tissue was trimmed into fragments of 6.5 x 6.5 x 5 mm3 from each fresh sample and then embedded in OCT using the operating instructions provided by 10x Genomics. The OCT-embedded tissues were then snap-frozen using isopentane pre-cooled with liquid nitrogen. Next, frozen sections of 4μm thickness were serially cut from the three tissue blocks. After Hematoxylin-eosin staining (H&E Staining Kit, Abcam, ab245880), the frozen sections were mounted on standard glass slides and reviewed by two expert pathologists to confirm the pathological diagnosis. The next continuous slides were used for RNA quality detection and processing sequence.

**Figure 1:**
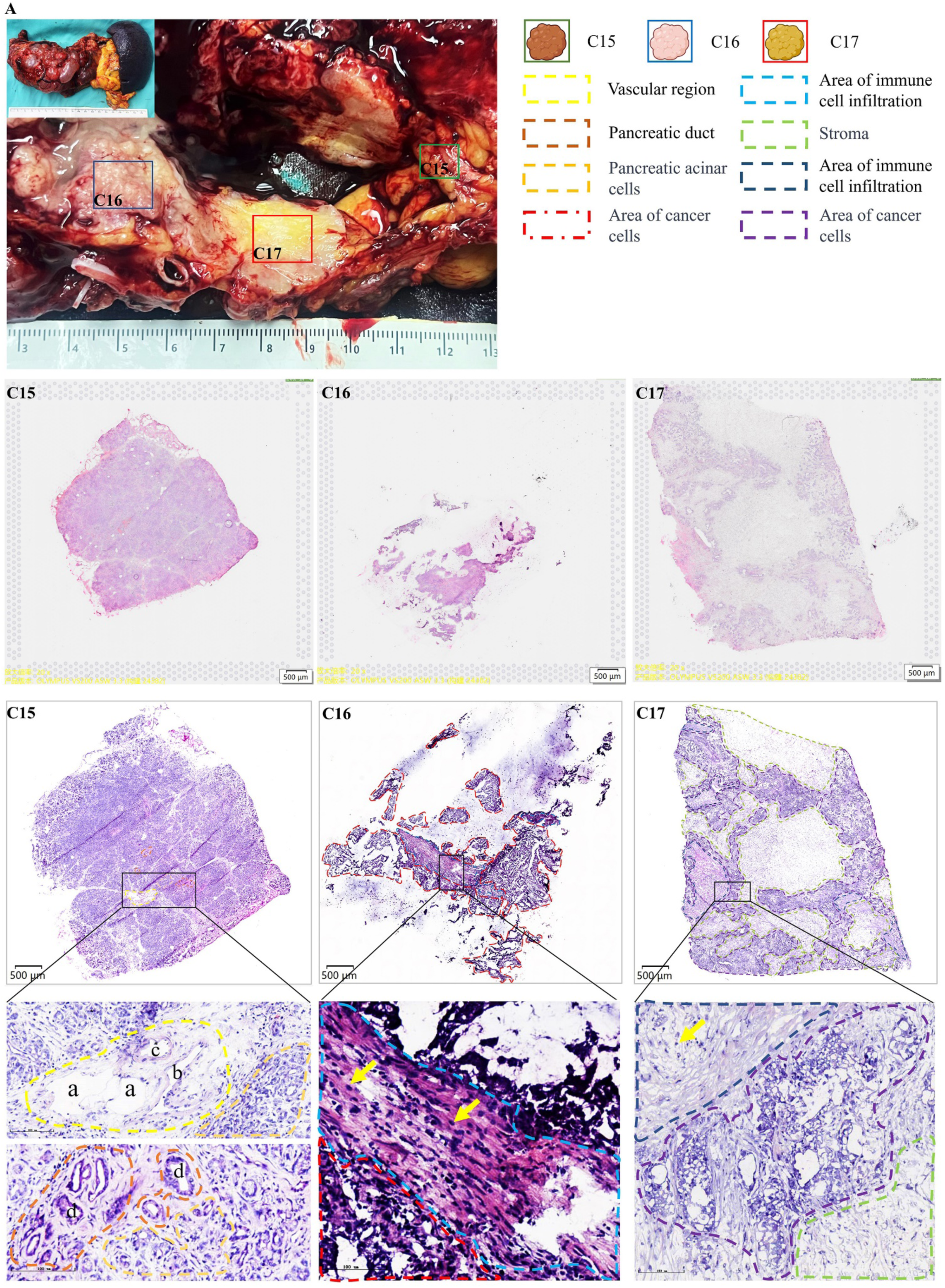
Sample collection, HE staining, and pathological annotation of pancreatic tumor. A. A 66-year-old female patient was admitted due to the “discovery of pancreatic cyst for 13 years, left upper abdominal distension for one month”. The preoperative diagnosis was multiple pancreatic cysts. “Laparoscopic pancreatic body and tail resection + splenic vein + portal vein repair surgery” was performed. The pathological results of the postoperative gross specimen showed pancreatic ductal papillary mucinous neoplasm with high-grade intraepithelial neoplasia, canceration, and low to medium differentiated adenocarcinoma formation. C15: Normal pancreatic tissue, annotated as vascular area, pancreatic ductal area, and acinar area. a. lymphatic vessel; b. vein; c. artery; d. pancreatic duct. C16: Two main areas were annotated, the immune infiltrated and tumor areas. The tumor area corresponds to the pancreatic ductal papillary mucinous neoplasm diagnosis with high-grade intraepithelial neoplasia. C17: Three main areas were annotated, immune infiltrated area, stroma, and tumor area. The tumor area corresponds to low to medium-differentiated adenocarcinoma. The arrow in the patient image points to a lymphocyte.

### RNA Quality and Permeabilization

After two pathology experts confirm that C15 is a normal pancreas, C16 contains high-grade IPMN, and C17 contains invasive PDAC, we cut 10 µm-thick cryosections for RNA quality detection and tissue permeabilization experiments (10x Genomics, Visium Spatial Tissue Optimization, Rev A). Specifically, we used the Trizol method to extract total RNA from cryosections. RNA concentration was detected using a Qubit fluorometer (ThermoFisher, Q33238), and RNA integrity number (RIN) was measured using the Agilent 2100 bioanalyzer (BiOptic Inc., Taiwan, China). Tissue permeabilization was performed according to the instructions provided by 10x Genomics. The optimal permeabilization time for those pancreas samples was 6 minutes.

### Spatial Transcriptomics (ST) Sequencing and Data Processing

Libraries for all tissue sections were generated following the 10x Genomics Visium library preparation protocol and sequenced on a NovaSeq 6000 System (Illumina) using a NovaSeq S4 Reagent Kit (200 cycles, 20027466, Illumina), at a sequencing depth of approximately 250–400 M read-pairs per sample. Sequencing was performed using the following read protocol: read 1, 28 cycles; i7 index read, 10 cycles; i5 index read, 10 cycles; read 2, 91 cycles. The raw FASTQ files of the 3 samples were directly run using Space Ranger (version 2.0.0, 10x Genomics) according to the official tutorial (https://support.10xgenomics.com/)and reads were mapped to the human reference genome (GRCh38-2020-A). A median of 3383, 2730, and 2124 genes was obtained per spot for C15, C16, and C17 respectively.

### Spots-by-spots Pathologist Annotation Using Loupe Browser

The pathologists He and Gu annotated C15, C16, and C17 spot-by-spot following a consensus workflow using Loupe Browser 6.2 (10x Genomics) referenced to the continuous HE slides (Figure S1). The lower coverage spots (tumor cells less than 20%) and missing spots (regions that did not cover tissue or scanning/sectioning artifacts that were impossible to recognize the cell type). The final consensus annotation dataset consisted of a total number of 4786 spots (1573 spots in C15 including normal acinar (1024), pancreatic duct (459), and pancreatic islet (90); 392 spots in C16 including immune infiltration (64) and high-grade IPMN (328); and 2821 spots in C17 including invasive PDAC (1091), immune rich region (77), and Stroma (1653).

### Spatial Transcriptomics Data Analysis and Visualization

Data analysis and visualization were carried out using the Seurat (version 4.1.1) R packages. The mitochondria proportion was measured using the PercentageFeatureSet() function. Counts were normalized using the SCTransform() function with parameters do.scale = T, vars.to.regress = “percent.mt” and the high variable genes were calculated using VariableFeatures() function. Then 3 ST data as well as the high variable genes of each sample were merged directly using normalized data. The dimension reduction performed by the RunPCA() function as assay = “SCT”. The FindNeighbors() and FindClusters() functions were used to cluster the ST spots, the parameters ues were dims = 1:30 and resolution = 1.5. To identify marker genes of different clusters, we used the FindAllMarkers() function as the following parameter: test.use = “wilcox”, logfc.threshold = 0.5, min.pct = 0.5. Based on the annotation of CellMarker http://xteam.xbio.top/CellMarker/ and their associated biological function, we defined a total of 8 clusters among 3 samples, including pancreatic duct, acinar, pancreatic islet, IPMN, immune, stroma, and PDAC. Then we perform differential expressed genes analysis among defined clusters using the FindAllMarkers() function as the following parameter: test.use = “wilcox”, logfc.threshold = 0.25, min.pct = 0.1 and identified the markers genes of each cluster as top 10 ave_log2FC genes with adj.P.value < 0.05. The SpatialFeaturePlot(), SpatialDimPlot(), and Vlnplot() functions were used to visualize the data. The DEG analysis between clusters was performed using the FindMarkers() function as adj.P.value < 0.05. The AddModuleScore() function was used to calculate the enrichment score of a specific geneset, 100 control features were selected from the same bin per analyzed feature. The ggplot2 (version 1.12.0) was used for visualizing the differential expressed genes (for PDAC compared with IPMN, we set the cutoff as |ave_log2FC| > 1; for all tumor compared with ductal, we set the cutoff |ave_log2FC| > 2). The enrichR and clusterProfiler package (version 4.2.2) were used for Hallmark enrichment analysis.

### Spatial Clone Inferring and Clone Tree Building

To explore the evolutionary relationship of HG IPMN and Invasive PDAC, we performed siCNV analysis in consensus annotations spots using the R package SpatialInferCNV (version 0.1.0). The Pancreatic Ductal cluster spots of C15 were used as a reference control. For the global visualization of gene-level siCNV events, the genomic positions file (generated from Homo_sapiens.GRCh38.107.gtf) was used. The function CreateInfercnvObject() was used with the following parameters: chr_exclude = c(“chrM”). The run() function was used with the following parameter values: analysis_mode = “subclusters”, cluster_by_groups = T, HMM = T, denoise = T, num_threads = 2. Then supervised siCNV analysis was performed using the R package inferCNV using the node identity file instead of the annotation file. The following inferCNV run parameters were used: cutoff = 0.1, num_threads = 10, cluster_by_groups = TRUE, denoise = TRUE, HMM = TRUE. To construct these trees, shared CNVs across clusters were identified, assuming that a CNV cannot be reversed once it occurs, and this indicates that the cells in these clusters share a common ancestry. Based on this logic, ancestral relationships among clusters were identified, and clone trees were constructed. The branch lengths between subclones were calculated as the additional CNVs in the descendant clone plus a constant. logarithm (base 2) of the number of additional CNVs in the descendant clone divided by 2. The circle diameter corresponding to the clone was calculated as 10 times the logarithm (base 2) of the number of spots and divided by 2. Then all the circles and branches were scaled during the visualization. The differentially expressed genes among clones were identified using the FindMarkers() function as p_val_adj < 0.05. The ComplexHeatmap (version 2.10.0) R package was used to visualize the annotations of the row of CNV heatmap. We followed the tutorial of NG-CHM (https://www.ngchm.net/Downloads/ngChmApp.html) to check the location of genes of interest.

### SpaCET for Cell Type Deconvolution

We use SpaCET (https://gitHub.com/data2intelligence/SpaCET) to infer cell identities from the tumor ST data according to the tutorial. The create.SpaCET.object.10X() function was used to create the SpaCET object and the spots were manually filtered to refer to pathologist annotation. The “PAAD” cancer type was selected as the gene pattern dictionary of copy number alterations to deconvolve the cell types of each spot (including malignant, immune, and stromal cells). We identified the TME boundary using SpaCET.identify.interface() function setting Malignant Cutoff = 0.8 and using the SpaCET.visualize.spatialFeature() function for visualization. Then the mean proportion of specific cell types within the interface of HG-IPMN and Invasive PDAC were compared and visualized using R package ggplot2 (version 1.0.12).

### Microenvironment Crosstalk Analysis

We used the CellChat (version 1.6.0) R package to study the different spatial communication of IPMN/PDAC with Immune and Stroma. Briefly, the normalized ST data were used to create a CellChat object, and the secretory autocrine/paracrine signaling database was analyzed. After processing data with default parameters, the computeCommunProbPathway() and aggregateNet() functions were used for inferring the intercellular communication network of each ligand-receptor pair (L-R pairs). The CellChat objects were merged using the mergeCellchat() function. The total number of interactions and strength between HG-IPMN and Invasive PDAC were compared and visualized by the compareInteractions() function. The sum of information flow for each signaling pathway was compared and visulized by the rankNet() function using parameters as cutoff.pvalue = 0.05. The L-R pairs communication probabilities were compared and visualized by netVisual_bubble() function setting comparison.

## Results

### Spatial Depiction of the Normal Pancreas, IPMN, and PDAC in the Same Patient and Their Unique Immune Microenvironment

We conducted hematoxylin and eosin (H&E) staining and spatial transcriptomics (ST) analysis on continuous tissue slides from a single patient. The slides included C15 (normal pancreas tissue from the tail), C16, and C17 (details provided in Figure 1A). Two senior pathologists annotated the various histological features on these slides, and the corresponding spatial spots were annotated using the Loupe Browser (Figure S1A). In slide C15, we classified three primary regions: the vascular, pancreatic ductal, and pancreatic acinar zones. In slide C16, we identified the immune infiltrate and tumor cell zones, while slide C17 revealed the stromal zone, the immune cell infiltrate zone, and the tumor cell zone (Figure 1). After performing quality control, which involved filtering out spots that did not cover any cells, we detected approximately 4,786 spots across the slides (C15: 1,573 spots; C16: 392 spots; C17: 2,821 spots). We then conducted principal component analysis (PCA) to reduce data dimensionality using shared variably expressed genes, followed by UMAP clustering.

The analysis revealed seven clusters, which were defined as Acinar, Ductal, and Islet in C15; IPMN, Stroma, and Immune in C16; and PDAC, IPMN, Stroma, and Immune in C17, based on differentially expressed genes (DEGs) across clusters. The spatial distribution of these clusters is shown in Figure 2D. Notably, the annotated spots were highly consistent with histological consensus annotation (Figure 2B). Interestingly, some spots in C17 clustered as IPMN (Figure 2C), suggesting transcriptomic heterogeneity in cancer and potential evolutionary processes.

**Figure 2.**
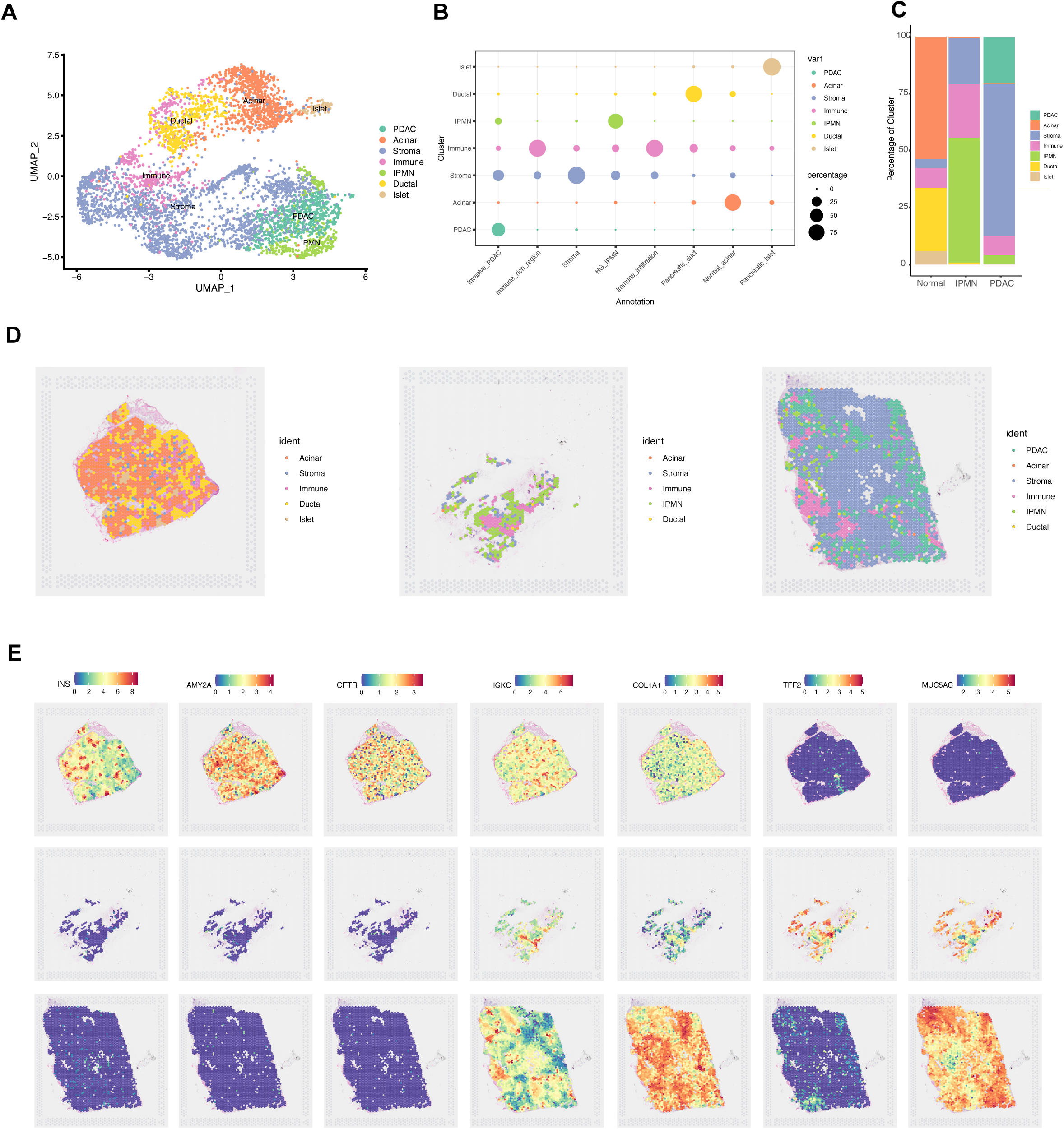
Spatial Clustering and Expression of Marker Genes. A. UMAP embedding of all clusters among three samples. B. Bubble plot showing the overlap between pathologist consensus annotations and spatial clusters. C. Stacked plot displaying the proportion of each cell type cluster across the three samples. D. Spatial maps illustrating the clusters in samples C15, C16, and C17. E. Spatial distribution of representative marker genes for each identified cluster.

Our findings identified marker genes for various regions: Islet (INS), normal pancreatic ductal cells (CFTR), Acinar (AMY2A), Immune (IGKC), Stroma (COL1A1), IPMN (TFF2), and PDAC (MUC5AC). The spatial expression patterns of these representative marker genes are displayed in Figure 2E. Notably, MUC1 and MUC5AC were expressed in the tumor regions of both IPMN and PDAC, whereas MUC2 expression was absent (Figure S1B). According to the WHO classification of IPMN subtypes^11^, the pathological subtype in this patient corresponds to the pancreaticobiliary type, consistent with the clinicopathological diagnosis. The molecular subtypes of PDAC are divided into several subtypes, with basal and classical subtypes being the most prominent^12,13,14^, the classical subtype is associated with better survival and drug response^15^, and IPMN-associated PDAC typically exhibits a classical transcriptional profile^16^. To further explore the molecular subtype in this patient, we calculated the subtype score on the ST data using Moffitt’s subtype signature genes. The results demonstrated that the tumor regions of HG-IPMN and Invasive PDAC predominantly enriched Moffitt’s classical score (Figure S1C). Thus, the tumor molecular subtype in this patient likely belongs to the classical subtype, with no significant intra-tumoral subtype heterogeneity observed.

### Dynamic Transcriptomic Changes During Tumor Progression

To analyze the changes in transcriptional profiles during tumor progression, we compared differentially expressed genes (DEGs) across the previously defined Pancreatic Ductal, IPMN, and PDAC regions. As illustrated in Figure 3A, we identified 34 and 89 genes that were upregulated exclusively in IPMN (marked in red) and PDAC (marked in blue), respectively, compared with Pancreatic Ductal regions (avg_log2FC > 2 and p.adj < 0.05), including IPMN markers gene MUC6, and PDAC classical markers (TFF2, AGR3, and TSPAN8). Moreover, we identified two critical DEG patterns: Peak and Climbing. In the Peak pattern, genes such as TFF2, MUC6, and AQP5 (Figure 3B), as well as transcription factors (TFs) ONECUT3, SOX9, and GATA6 (Figure S2A and B), were upregulated in IPMN followed by downregulation in PDAC. This suggests that these genes may contribute to dysregulation of the pancreatic duct but not to malignant transformation. In contrast, the Climbing pattern showed that the expression of KRT17, AREG, and AGR3 (Figure 3B), as well as TFs GATA3, ELF3, KLF4, and HMGA1 (Figure S2A and B), increased progressively as the tumor advanced. This pattern indicates their role in promoting malignancy in the pancreatic duct and rendering cells more susceptible to malignant transformation.

**Figure 3.**
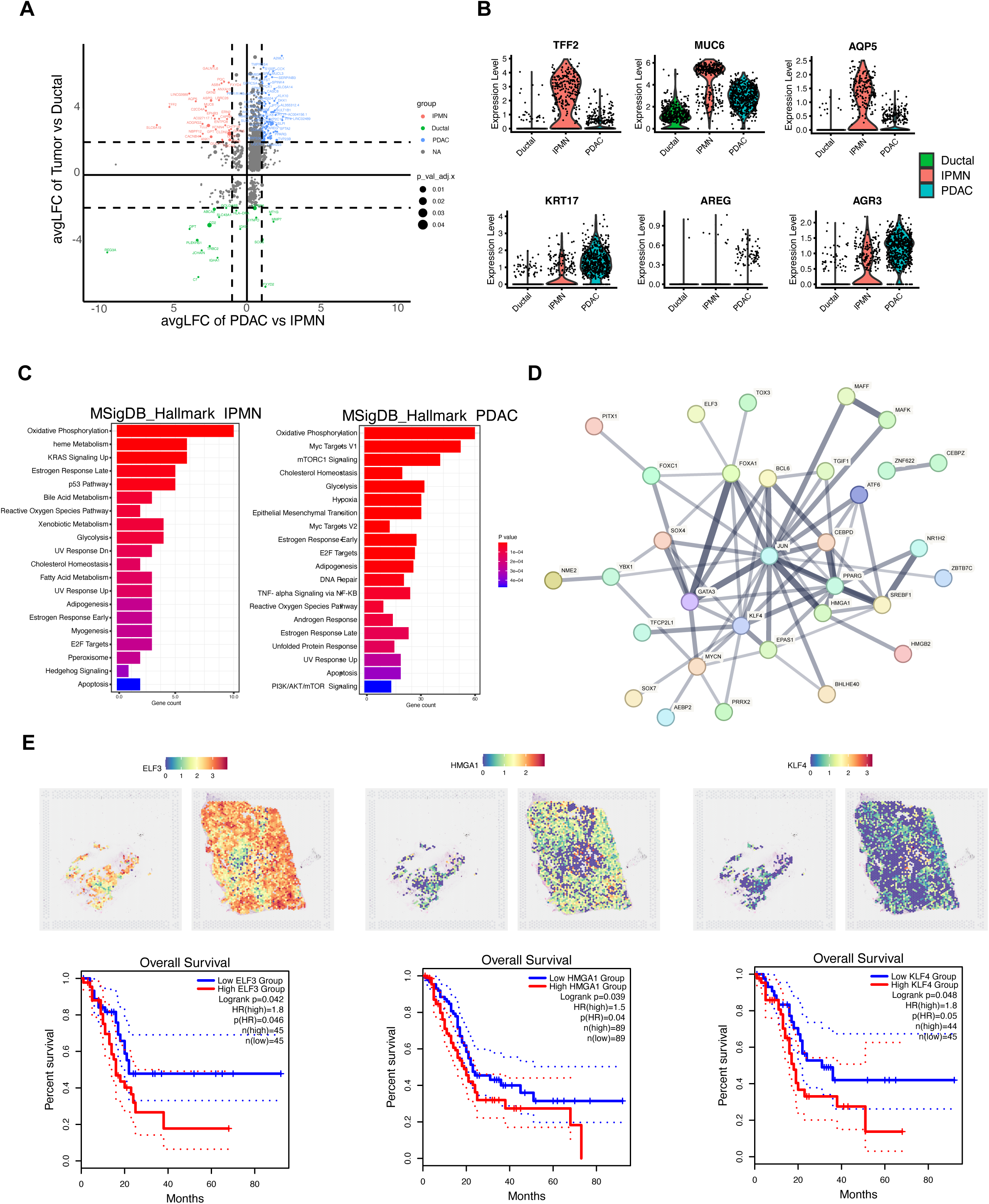
Transcriptomic Changes During Tumor Progression. A. Scatter plot illustrating differential gene expression across IPMN, PDAC, and Ductal regions. Representative genes exclusively up-regulated in each group are colored. B. Violin plot showing representative genes that are specifically up-regulated in IPMN (peak) or PDAC (climbing). C. Bar plot depicting HALLMARK pathway enrichment of differentially expressed genes in each cluster. D. PPI network analysis of transcription factors up-regulated in PDAC. E. Spatial expression and overall survival data (from TCGA) for representative transcription factors.

Next, we performed the enrichment analysis of hallmark pathways on IPMN, and PDAC up-regulated genes. Our findings revealed multiple pathways involved in tumor progression. Oxidative phosphorylation emerged as a critical pathway in both IPMN and PDAC, while glycolysis and cholesterol homeostasis were significant in either IPMN or PDAC. Notably, metabolic-related pathways were enriched in IPMN, including heme metabolism, bile acid metabolism, and fatty acid metabolism. In contrast, PDAC regions showed significant enrichment in oncogenic signaling pathways, such as MYC and mTORC1, as well as the EMT and hypoxia pathways, which are known to play crucial roles in tumorigenesis and progression (Figure 3C, Supplementary Figure S2C). These findings underscore the role of these pathways in the eventual transformation of HG-IPMN to invasive PDAC.

Subsequently, we performed a protein-protein interaction (PPI) network analysis on the set of TFs exclusively upregulated in the PDAC cluster. This analysis revealed that TFs such as JUN, GATA3, FOXA1, KLF4, and HMGA1 acted as core regulators of tumor progression (Figure 3D). Among these TFs, higher expression levels of ELF3, HMGA1, and KLF4 were not only observed in the PDAC cluster compared with the IPMN cluster but were also associated with a worse prognosis in TCGA PDAC patient samples (Figure 3E). These results provide valuable insights and directions for further in-depth studies on the mechanisms underlying PDAC development.

### Copy Number Variation and Clonal Evolution Analysis of High-Grade IPMN and Invasive PDAC

Tumorigenesis and progression are often associated with dynamic gene expression changes and spatiotemporal heterogeneity at different stages^17^. To investigate the progression of IPMN, we calculated chromosomal copy number variation (CNV) in tumor cell spots of HG IPMN and Invasive PDAC using SpatialInferCNV (the Ductal cluster was used as the benign reference). Our findings revealed significant tumor clonal heterogeneity in the PDAC sample, while the IPMN sample exhibited only a single homogeneous clone. As illustrated in Figure 3A, invasive PDAC shared a similar profile of CNV events with HG IPMN, characterized by gains in chromosomes 1, 3, 7, 8, 9, and 17 and losses in chromosomes 1, 4, 5, 6, 7, 10, 13, 14, 16, 20. This suggests that invasive PDAC may originate from the same clone as HG IPMN. Regarding oncogenes, both HG-IPMN and invasive PDAC harbored amplifications of MUC1 (chr1), PIK3CA (chr3), EGFR (chr7), and MYC (chr8), along with losses of tumor suppressor genes such as EPHA2 and PTEN. Additionally, both exhibited ELF3 (chr1) gain, consistent with our DEG analysis, and the transcription factor KLF4 (chr9) appeared to undergo copy number gain as the tumor progressed (Figure 4A).

**Figure 4.**
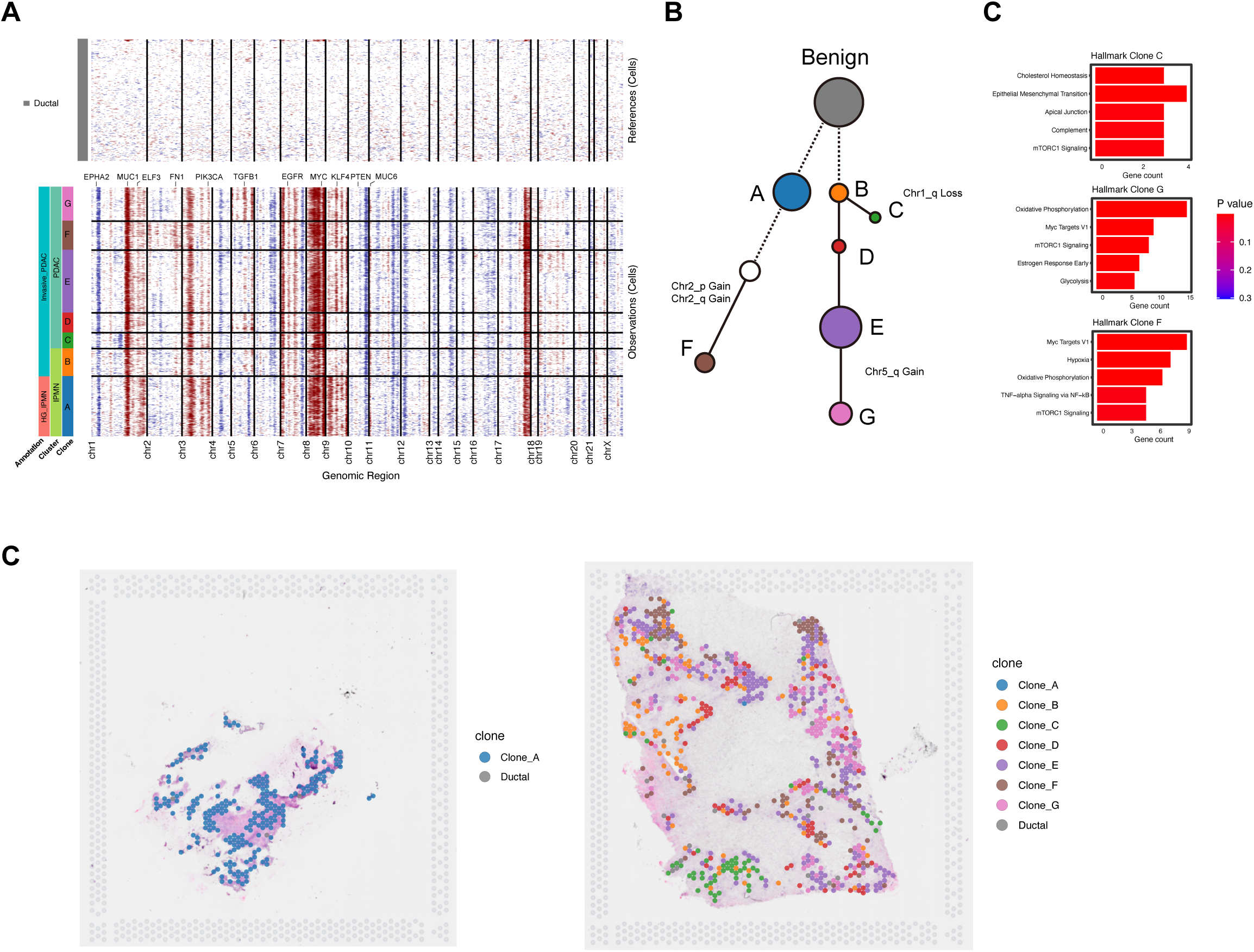
Copy Number Variation (CNV) Analysis in Pancreatic Ductal, HG IPMN, and Invasive PDAC. A. Hierarchical clustering heatmap displaying CNV events across all spots in HG IPMN and Invasive PDAC, using pancreatic duct spots as a normal reference. Genes of interest with significant CNV changes are annotated. B. Phylogenetic clone tree of pancreatic duct, HG IPMN, and Invasive PDAC. Circle size represents the number of spots in each clone, and line length indicates the normalized increment of CNV events between parent clones. Grey circles denote hypothetical clones. C. Top 5 functional enrichment pathways identified in each terminal clone. D. Spatial distribution of all clones in HG-IPMN and Invasive PDAC.

Using an unsupervised approach for CNV clustering, invasive PDAC was divided into seven distinct subclones (Clones A-G) (Figure 4A). Interestingly, Clone B corresponded to the IPMN cluster within the invasive PDAC sample and had minimal CNV events. The seven subclones of invasive PDAC demonstrated distinct spatial locations and heterogeneity, with significant changes in certain chromosomes (Figure 4C). For instance, Clone F exhibited amplification on chromosome 2, including FN1 gene gain, while Clones D, E, and G showed amplification on chromosome 5 compared with others (Figure 4A).

As the number of CNV events increased from Clones A to G, and considering the irreversibility of CNV variants, we constructed a clonal evolutionary tree mapping the spatial reconstruction from HG IPMN to invasive PDAC. As shown in Figure 4B, Clone F of invasive PDAC is inferred to have evolved from Clone A of HG IPMN, primarily through a gain in chromosome 2q, which includes the FN1 gene. Additionally, Clones D, E, and G were inferred to have evolved from Clone B, showing CNV gain of chromosome 5q with TGFB1 gene gain. To understand the distinct functions of each terminal clone, we performed enrichment analysis on Clones F, G, and C. The results showed that Clone F was significantly enriched in hypoxia-related genes, Clone G had higher activation of oxidative phosphorylation, and Clone C was enriched with EMT-related genes (Figure 4D. The above evidence supports that Invasive PDAC demonstrated relatively more malignant and intra-tumoral heterogeneity, with different functional clones accumulating CNV events compared with HG IPMN.

### mCAF Exhibits a Closer Interaction with the Progression from HG IPMN to Invasive PDAC

Cancer-associated fibroblasts (CAFs) are one of the most abundant stromal cell types in the tumor microenvironment (TME). They play a critical role in reshaping the extracellular matrix (ECM) and promoting tumor progression by secreting various growth factors, chemokines, and cytokines, as well as through interactions with tumor cells and immune cells^18^. Using SpaCET to decompose cell lineages by integrating gene expression patterns of PAAD from TCGA, we inferred the abundance of malignant cells, CAFs, endothelial cells, and various immune cells for each spatial spot. To explore the differences in the adjacent TME between high-grade (HG) IPMN and invasive PDAC, we set the boundary of the tumor area as spots where the proportion of malignant cells exceeded 80% (excluding spots containing 100% malignant cells), covering the majority of previously annotated IPMN and PDAC regions (Figure 5A). We found that the proportion of CAFs within HG IPMN was 20.21%, which increased to 40.13% within invasive PDAC (Figure 5B). We calculated the module scores for mCAF and iCAF signature genes to investigate the differences in their spatial distribution across each tumor region. The results showed greater enrichment of both mCAF and iCAF in invasive PDAC compared to HG IPMN, particularly for mCAF (Figure 5B). Moreover, mCAFs were more enriched than iCAFs in both HG IPMN and invasive PDAC (both P < 2.22e-16, respectively; Figure 5C). In non-tumor areas of HG IPMN and invasive PDAC, mCAF enrichment was also predominant over iCAF (Figure S2D).

**Figure 5.**
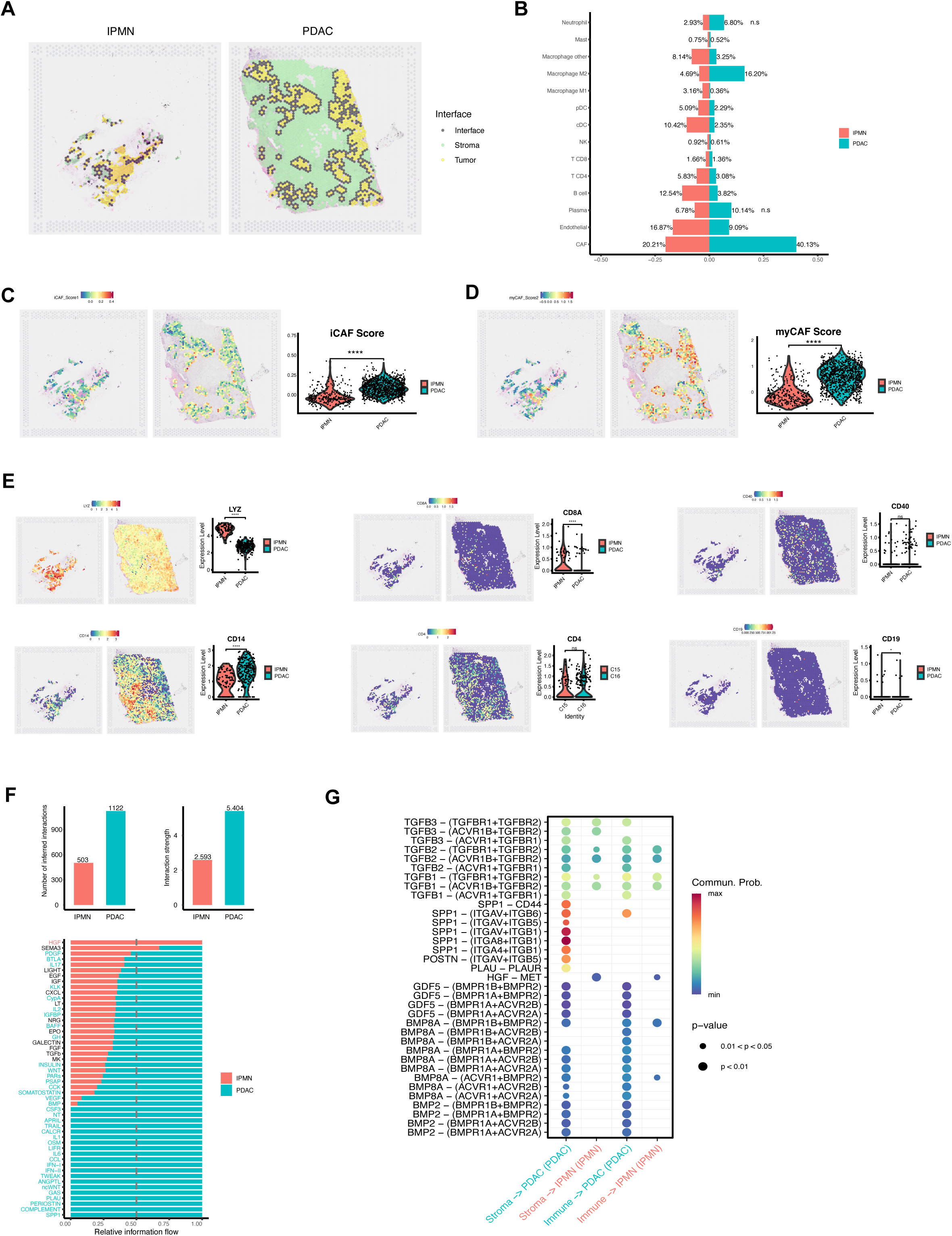
Spatial Heterogeneity of the Microenvironment and Immune Infiltration Between HG IPMN and Invasive PDAC. A. Inferred tumor regions (yellow spots) defined by malignant cell proportion > 80%, with neighboring spots defined as interface boundaries (black spots). B. Stacked plot showing the proportions of different cell types within the two tumor regions. C. Spatial enrichment map of iCAF signature scores within tumor regions (excluding spots with 100% malignant cells). Violin plot comparing scores between HG-IPMN and Invasive PDAC. D. Spatial enrichment map of mCAF signature scores within tumor regions (excluding spots with 100% malignant cells). Violin plot comparing scores between HG-IPMN and Invasive PDAC. E. Relative expression and spatial distribution of LYZ, CD14, CD4, CD8A, CD19, and CD40 in each tumor region. F. Bar plot illustrating the total number of ligand-receptor pair interactions between HG-IPMN/Invasive PDAC and their respective Immune/Stroma clusters. G. Stacked plot ranking significant information flow differences between HG-IPMN and Invasive PDAC with their respective Immune/Stroma clusters (red/blue labels indicate significance). H. Ligand-receptor interactions from the Immune/Stroma cluster to the tumor cluster. The y-axis lists ligand and receptor names, while the x-axis represents the interacted regions. Dot color reflects communication probabilities, and dot size represents computed p-values.

### Immune Landscape Changes and Intercellular Signaling Differences Between High-Grade IPMN and Invasive PDAC

Myeloid cells and T cells are critical components of the immune landscape within the cancer ecosystem, playing vital roles in tumor progression through both direct and indirect mechanisms^19^. Based on marker expression and spatial feature map percentages, myeloid cells marked by LYZ and monocytes marked by CD14 were more abundant than T cells marked by CD4A and CD8 in both HG-IPMN and invasive PDAC across the sections and within their tumor regions (Figure 5D). Notably, compared to invasive PDAC (malignant > 0.8), IPMN had a higher proportion of CD4+ T cells (5.38% vs. 3.08%), B cells (12.54% vs. 3.82%), type I interferon (IFN-I) producing plasmacytoid dendritic cells (pDCs) (5.09% vs. 2.29%), and conventional dendritic cells (cDCs) (10.42% vs. 2.35%). These immune cells are known to effectively activate CD4+ and CD8+ T cells and induce cytotoxic T lymphocytes (CTLs) production, leading to anti-tumor effects (Figure 5B)^20^. In contrast, PDAC was predominantly characterized by a higher infiltration of M2 macrophages (16.20% vs. 4.69%) (Figure 5A), which are associated with promoting tumor growth, metastasis, and inhibiting other immune cells, thereby preventing the immune system from mounting an effective anti-tumor response^21,22,23^. Additionally, PDAC exhibited a reduction in CD4+ T cells, CD8+ T cells, NK cells, and B cells—all of which have anti-tumor effects—indicating that the tumor environment had transitioned into a “cold tumor” state with immune suppression as HG-IPMN progressed to invasive PDAC^24^.

We subsequently employed CellChat to address the differences in crosstalk between HG-IPMN/Invasive PDAC with Immune/Stroma regions. Firstly, we compared the information flow^25^ for each secretory signaling pathway between the two groups. We observed that the interactions between PDAC and the immune/stroma regions were more numerous and stronger than those in IPMN. The signaling probability of receptor–ligand interactions revealed that HGF was uniquely present in HG-IPMN but not in invasive PDAC. Conversely, tumor-promoting pathways such as SPP1, TGFb1, POSTN, and ncWNT signaling were widely enhanced in PDAC. In particular, the stroma cluster in invasive PDAC was the predominant sender of interactions with the PDAC cluster (Figure 5G). These findings indicate that the microenvironment in PDAC acquires a tumor-promoting and immune-evasive phenotype.

## Discussion

In this study, we comprehensively explore the spatial and molecular landscape of normal pancreas, high-grade intraductal papillary mucinous neoplasm (HG-IPMN), and invasive pancreatic ductal adenocarcinoma (PDAC) within the same patient. By integrating spatial transcriptomics and multi-analytical approaches, we reveal crucial insights into the transition from IPMN to invasive PDAC. Our analysis demonstrates significant transcriptomic heterogeneity across normal, IPMN, and PDAC tissues, identifying key transcription factors, such as ONECUT3, SOX9, and GATA6, that contribute to dysregulation in IPMN but not malignant transformation, while GATA3, ELF3, KLF4, and HMGA1 progressively increase during tumor progression, driving malignancy. Notably, oxidative phosphorylation emerges as a pathway common to both IPMN and PDAC, with oncogenic pathways such as MYC, mTORC1, EMT, and hypoxia being significantly activated during malignant transformation. Copy number variation (CNV) analysis further supports the clonal evolution of PDAC from HG-IPMN, with invasive PDAC exhibiting significant spatial heterogeneity compared to the more homogeneous IPMN. Importantly, our spatial analysis highlights the role of myofibroblastic cancer-associated fibroblasts (mCAFs), which are more enriched in invasive PDAC and closely interact with tumor cells, driving progression. The immune landscape analysis reveals a shift from a relatively immune-active environment in HG-IPMN, characterized by CD4+ T-cells and dendritic cells, to a highly immune-suppressive “cold” tumor microenvironment (TME) in invasive PDAC, dominated by M2 macrophages. These findings are further reinforced by intercellular signaling analysis, which shows enhanced tumor-promoting pathways in PDAC, indicating a shift towards an immune-evasive phenotype.

Transcript factors and pathways that may play critical roles in the natural progression of pancreatic cancer. Our findings showed that several genes, including MYC, ELF3, and KLF4 exhibited copy number gains in HG IPMN and Invasive PDAC. Numerous reports have documented that MYC overexpression can lead to tumorigenesis^26^. MYC activation drives cancer progression through two mechanisms: acquiring cancer hallmarks intrinsic to tumor cells or changes in the tumor microenvironment (TME) and anticancer immune response^27,28^. ELF3 belongs to the epithelium-specific ETS (ESE) transcription factor family. Genetic alterations and imbalanced expression of ESE transcription factors are associated with the progression of several epithelial cancers, such as lung, breast, and colorectal cancer^29^. Embryonic deletion of KLF4 leads to alterations in skin permeability and postnatal death, apart from that, it is also reported to play a regulator role in malignancy^30^. KLF4 protein was increased in early lesions of PDAC and promoted the formation of pre-cancerous pancreatic lesions in the mouse model^31,32^. Ganguly, et al. discovered a strong positive correlation of Muc5ac with Klf4 in the PanIN lesions of KC mouse pancreas, reinforcing the crucial involvement of KLF4 in bolstering the CSC-associated tumorigenic properties and leading to pancreatic cancer onset and progression^33^. We know that tumorigenesis also involves complex cellular signaling pathways. Given the HALLMARK enrichment analysis of signaling pathways, we found gradual enrichment scores for multiple pathways involved in tumor metabolism, proliferation, signaling transduction, and immunity et al. These pathways are interconnected in a network and mutually promote the occurrence and development of tumors. For instance, glucose-dependent metabolism, particularly aerobic glycolysis, plays a crucial role in the immune function of CD8^+^ effector T cells and the IFN-γ production of CD4+ T cells^34,35^. In contrast, to CD4^+^ effector T cells (Th1, Th2, and Th17) and M1 macrophages, Tregs and M2 macrophages are primarily driven by lipid oxidation and rely less on glycolysis^36^. Our analysis reveals that aggressive pancreatic invasive PDAC is enriched with high-scoring metabolic pathways (oxidative phosphorylation, glycolysis, adipogenesis) while being accompanied by a low proportion of CD4^+^ T/CD8^+^ T immune cells and a high proportion of M2 macrophages.

In the natural progression of tumors, tumor cells gradually accumulate molecular events that confer a survival advantage in natural selection, including copy number alterations. Such copy number alterations are closely associated with tumor heterogeneity^37^. We inferred genome-wide information in each spot, which facilitated data-driven clone generation in a tissue-wide fashion at high resolution^38^. We discovered that the subclones of Invasive PDAC had evolved from the HG-IPMN. There was a high level of intra-tumoral heterogeneity among the subclones of Invasive PDAC. In our study, both HG-IPMN and invasive PDAC harbored amplifications of MUC1 (chr1), PIK3CA (chr3), EGFR (chr7), and MYC (chr8), along with losses of tumor suppressor genes such as EPHA2 and PTEN. Those findings in line with previous WGS analysis in IPMN of mostly mixed phenotypes revealed recurrent gains in chromosomes 3, 7, and 8^6,39^ (ref). Consistent with previous ST inferCNV analysis on the prostate, we found that clone B harbored fewer CNV events than other clones, which were also defined as the IPMN cluster based on the transcriptomic features. Taking advantage of the spatial information, we could identify clones that are hard to distinguish by histological morphology or even random sampling of single cells^38^. Notably, our findings reveal not only spatial heterogeneity of invasive PDAC but also demonstrate that the clones work as distinct functional during progression. This observation underscores the dynamic nature of tumors and the fact that cancer typically becomes increasingly heterogeneous as the disease progresses^40^.

PDAC is characterized by the abundant presence of non-malignant stromal cells, accounting for up to 90% of the tumor volume^41^. These stromal cells support cancer cell proliferation, survival, and invasion^42^. CAFs are a heterogeneous group among the stromal cells, including myofibroblastic CAFs (mCAF), inflammatory CAFs, and antigen-presenting subtypes^43^. The former is adjacent to tumor cells, expresses α-smooth muscle actin (αSMA), and secretes factors that stimulate tumor growth, cell survival, and metastasis; the latter is more distantly distributed throughout the tumor, low expresses α-smooth muscle actin and high secrets cytokines such as IL-6 and IL-11, which interact with tumor cells^44^. By deconvoluting the cell type proportion of each spatial spot, we can identify the most adjacent TME and explore the peritumoral crosstalks among multi-cell types. Our findings revealed that compared to iCAF, mCAF had a higher proportion around the tumor region, and played a more intimate role in the tumor. It has been reported that CAFs are dynamic and can exhibit different phenotypes depending on their spatial and biochemical niches in the PDAC microenvironment^44^. Tumor-secreted TGF-β can transform adjacent stromal cells into mCAF^45^, producing an extracellular matrix (ECM) around the tumor. However, this mechanical barrier can hinder proper blood vessel formation, limiting chemotherapy exposure and impeding immune cell infiltration^46^. We also evaluated the immune cell infiltration and differences in cell-cell communication between HG-IPMN and Invasive PDAC. Our results showed that in Invasive PDAC, the proportion of anti-tumor immune cells (CD4^+^ and CD8^+^ T cells) decreased, while the proportion of pro-tumor M2 macrophages increased compared to HG-IPMN.

The development and progression of pancreatic cancer are strongly influenced by tumor-associated inflammation within and surrounding the tumor^47,48,49^. Using spatial transcriptomics to analyze the proportions of immune cells in HG-IPMN and Invasive PDAC, we found that HG IPMN contains anti-tumor immune components, which appear to be gradually lost during tumor progression, accompanied by the accumulation of immunosuppressive cells. Our results further illustrate that, compared to “hot” immune tumors such as melanoma with high neoantigen loads and robust T cell infiltration, HG-IPMN progresses towards an “immunologically cold” tumor^50^. The main factor driving this result is that tumor cells shape an immunosuppressive TME through multiple pathways^50^, such as the mechanical barrier caused by the deposition of extracellular matrix ECM induced by mCAFs mentioned before. In addition, IGFBP2 can polarize macrophages towards the M2 state^51^. Experiments in mouse models have shown that IGFBP2 promotes tumor growth and facilitates an immune-suppressive microenvironment. We also studied the differences in cell-cell communication in HG IPMN and Invasive PDAC, the TGF-β pathway was activated between tumor and stroma in invasive PDAC. It is worth noting that the communication signaling pathway involving SPP1 is also switched on in invasive PDAC. SPP1 is a chemokine-like sialic acid-rich glycoprotein overexpressed in various cancers^52,53^. Consistent with previous findings, SPP1 upregulation is associated with tumor progression and a poorer prognosis^54^.

Despite these findings, several limitations should be acknowledged. First, the observed molecular mechanisms and cell-cell interactions are needed to further validate by in vitro and in vivo functional experiments. Second, the small sample size is a limitation due to the rarity of matched samples from the same patient, which may impact the generalizability of our findings. Integrating more samples and leveraging single-cell RNA sequencing (scRNA-seq) data could enhance the robustness of our results and provide deeper insights into IPMN-PDAC progression. Additionally, the spatial transcriptomics (ST) approach has inherent resolution limitations, which restrict the identification of fine-scale cellular heterogeneity compared to scRNA-seq. Integrating ST with scRNA-seq could help overcome these limitations, offering a more comprehensive view of the spatial and molecular dynamics at play.

In conclusion, our study provides an in-depth spatial and molecular characterization of IPMN malignant transformation and the progression to invasive PDAC. We identify key transcriptional programs, CNV-driven clonal heterogeneity, and distinct TME features that contribute to this transition. These findings highlight the complexity of pancreatic tumor evolution and underscore the need for early diagnostic biomarkers and therapeutic strategies that target both tumor cells and their microenvironment. Integrating spatial transcriptomics with single-cell data could pave the way for a more comprehensive understanding of pancreatic cancer biology, potentially leading to better therapeutic interventions. Future research should focus on validating these findings in larger cohorts and further exploring the role of mCAFs and immune modulation in IPMN-associated PDAC progression.

## Ethics approval and consent to participate

All experimental procedures related to sample collection and operation were approved by the Biomedical Research Ethics Committee of West China Hospital of Sichuan University, with Research Ethics Board approval following the Declaration of Helsinki. Written informed consent was obtained from this study participants.

## Availability of data and materials

The datasets used and/or analyzed during the current study are available from the corresponding author upon reasonable request.

## Authors’ contributions

SRP, QXC, and ZXC: Conception and design of the work; analysis and interpretation of data; drafting the work. DH and XFG: Conception of the work; acquisition, analysis. YC, KC, and MLY: Conception and design of the work; acquisition of data. JL, HC, and PG: Conception and design of the work; acquisition and analysis of data. XG, YQC, XW, YBL, and MZ: Acquisition, analysis, and interpretation of data. LWM, QX, PPX, and JZ: Analysis, and interpretation of data. BP and ZW: Conception and design of the work; acquisition, analysis, and interpretation of data; revising the work and approving the manuscript.

## Acknowledgments

We are most grateful to the Core Facility of West China Hospital for their technical support. We thank Li Li, Fei Chen, and Chunjuan Bao (Institute of Clinical Pathology, West China Hospital) for their efforts in performing frozen sectioning on our fresh tissues.

## Supplementary Figure Legend

**Figure S1.**
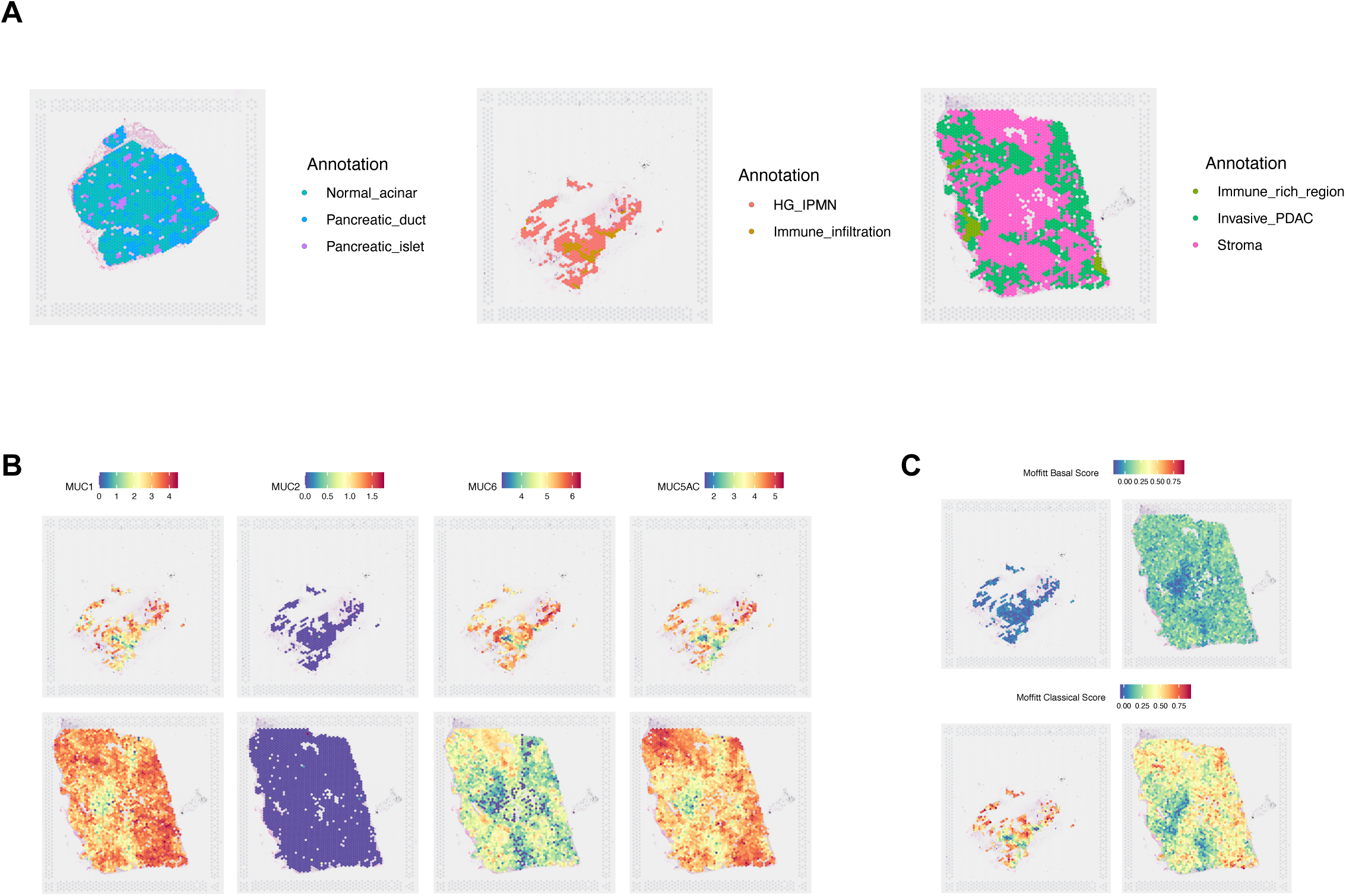
Transcriptomic Features in IPMN and PDAC. A. Spatial map of pathologist annotations for IPMN and PDAC. B. Distribution of representative marker genes (MUC1, MUC2, MUC6, and MUC5AC) relevant to IPMN pathological subtype classification. C. Spatial enrichment scores of Moffitt’s Basal/Classical signature in tumor regions. D. Spatial enrichment of representative HALLMARK gene sets in IPMN and PDAC.

**Figure S2.**
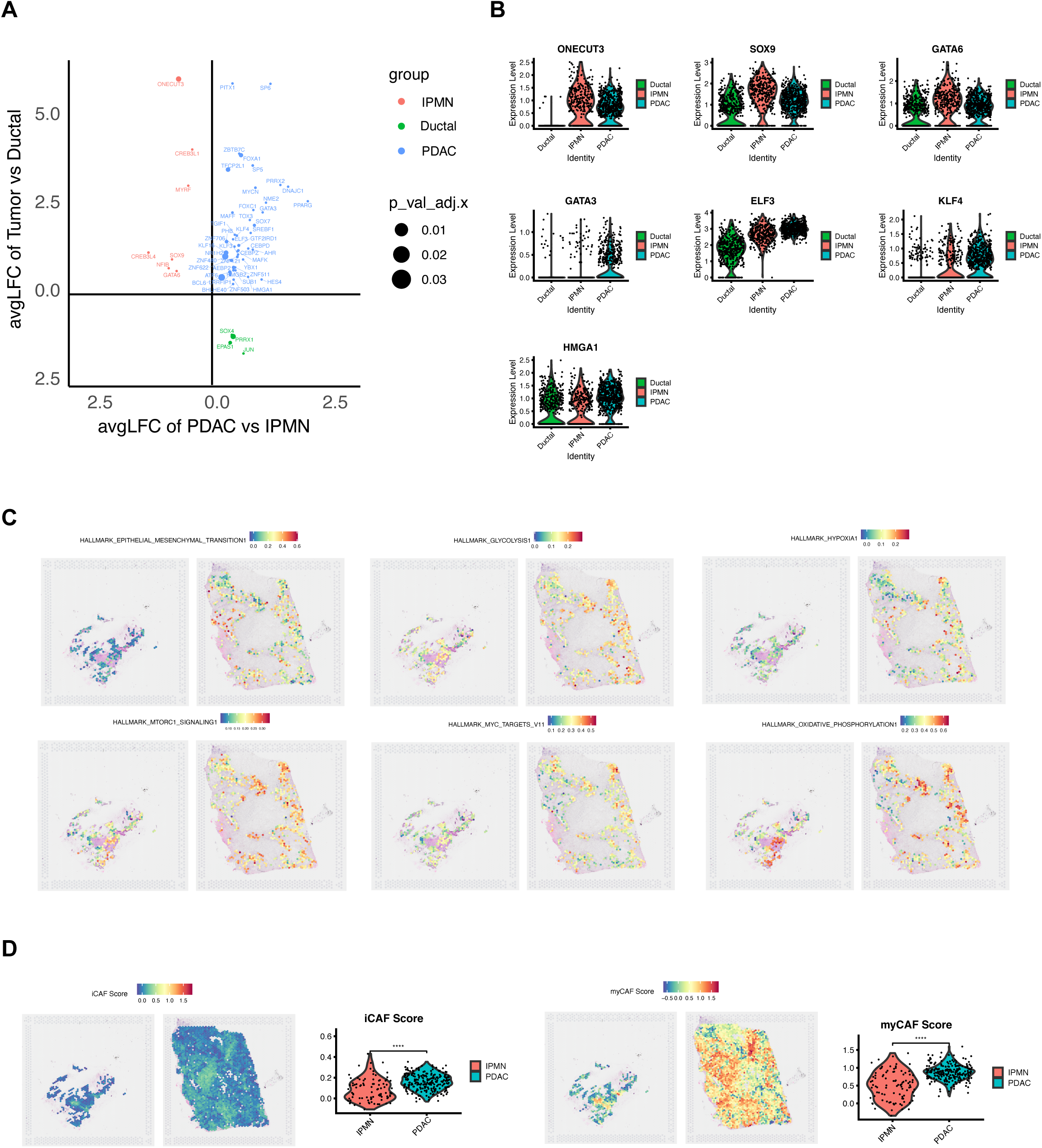
Spatial Transcriptomic Atlas Differences Between IPMN and PDAC. A. Scatter plot showing differential expression of transcription factors across IPMN, PDAC, and Ductal regions. Representative genes up-regulated in each group are highlighted. B. Violin plot depicting expression levels of representative transcription factors across IPMN, PDAC, and Ductal clusters. C. Spatial enrichment score map for representative HALLMARK pathways, showing data exclusively from tumor regions. D. Spatial distribution of iCAF and mCAF signature scores in IPMN and PDAC, with a violin plot comparing scores within the Stroma cluster regions only.

## References

1. Rahib L, et al. Estimated projection of US cancer incidence and death to 2040. JAMA Netw Open 4(4) (2021), e214708.

2. Basturk O, et al. A Revised Classification System and Recommendations From the Baltimore Consensus Meeting for Neoplastic Precursor Lesions in the Pancreas. Am J Surg Pathol 39(12) (2015), 1730–1741.

3. Wu J, et al. Recurrent GNAS mutations define an unexpected pathway for pancreatic cyst development. Sci Transl Med. 3(92) (2011), 92ra66

4. Tanaka M. Intraductal papillary mucinous neoplasm of the pancreas: diagnosis and treatment. Pancreas. 28(3) (2004), 282–8

5. Furukawa T, et al. Whole-exome sequencing uncovers frequent GNAS mutations in intraductal papillary mucinous neoplasms of the pancreas. Sci Rep. 1 (161) (2011).

6. Fischer CG, et al. Intraductal Papillary Mucinous Neoplasms Arise From Multiple Independent Clones, Each With Distinct Mutations. Gastroenterology. 157(4) (2019), 1123–1137.

7. Bernard V, et al. Single-Cell Transcriptomics of Pancreatic Cancer Precursors Demonstrates Epithelial and Microenvironmental Heterogeneity as an Early Event in Neoplastic Progression. Clin Cancer Res. 25(7) (2019), 2194–2205.

8. 10x Genomics sequences in situ. Nat Biotechnol. 38 (11) (2020), 1222

9. Sun L, et al. Single-cell and spatial dissection of precancerous lesions underlying the initiation process of oral squamous cell carcinoma. Cell Discov 9 (28) (2023).

10. Lv J, et al. Spatial transcriptomics reveals gene expression characteristics in invasive micropapillary carcinoma of the breast. Cell Death Dis 12 (1095) (2021).

11. Nagtegaal ID, et al. Board WHOCoTE: The 2019 WHO classification of tumours of the digestive system. Histopathology 76 (2020), 182–188.

12. Bailey P, et al. Genomic analyses identify molecular subtypes of pancreatic cancer. Nature 531 (2016), 47–52.

13. Moffitt RA, et al. Virtual microdissection identifies distinct tumor- and stroma-specific subtypes of pancreatic ductal adenocarcinoma. Nat Genet 47 (2015), 1168–1178.

14. Collisson EA, et al. Subtypes of pancreatic ductal adenocarcinoma and their differing responses to therapy. Nat Med 17 (2011), 500–503.

15. Krieger TG, et al. Single-cell analysis of patient-derived PDAC organoids reveals cell state heterogeneity and a conserved developmental hierarchy. Nat Commun 12 (2021), 5826.

16. Collisson EA, et al. Molecular subtypes of pancreatic cancer. Nat Rev Gastroenterol Hepatol 16 (2019), 207–220.

17. Dagogo-Jack I, et al. Tumour heterogeneity and resistance to cancer therapies. Nat Rev Clin Oncol 15 (2018), 81–94.

18. Chen X, et al. Turning foes to friends: targeting cancer-associated fibroblasts. Nat Rev Drug Discov 18 (2019), 99–115.

19. Galon J, et al. Tumor Immunology and Tumor Evolution: Intertwined Histories. Immunity 52 (2020), 55–81.

20. Collin M, et al. Human dendritic cell subsets: an update. Immunology 154 (2018), 3–20.

21. Yang Q, et al. The role of tumor-associated macrophages (TAMs) in tumor progression and relevant advance in targeted therapy. Acta Pharm Sin B 10 (2020), 2156–2170.

22. Jetten N, et al. Anti-inflammatory M2, but not pro-inflammatory M1 macrophages promote angiogenesis in vivo. Angiogenesis 17 (2014), 109–118.

23. Mills CD, et al. A Breakthrough: Macrophage-Directed Cancer Immunotherapy. Cancer Res 76 (2016), 513–516.

24. Bonaventura P, et al. Cold Tumors: A Therapeutic Challenge for Immunotherapy. Front Immunol 10 (2019), 168.

25. Jin S, et al. Inference and analysis of cell-cell communication using CellChat. Nat Commun 12 (2021), 1088.

26. Dang CV. MYC on the path to cancer. Cell 149 (2012), 22–35.

27. Casey SC, et al. MYC: Master Regulator of Immune Privilege. Trends Immunol 38 (2017), 298–305.

28. Chappell J, et al. Roles for MYC in the establishment and maintenance of pluripotency. Cold Spring Harb Perspect Med 3 (2013), a014381.

29. Luk IY, et al. ELF3, ELF5, EHF and SPDEF Transcription Factors in Tissue Homeostasis and Cancer. Molecules 23 (2018).

30. He, Z, et al. KLF4 transcription factor in tumorigenesis. Cell Death Discov. 9, 118 (2023).

31. Wei D, et al. KLF4 Is Essential for Induction of Cellular Identity Change and Acinar-to-Ductal Reprogramming during Early Pancreatic Carcinogenesis. Cancer Cell. 29 (2016), 324–38.

32. Prasad NB, et al. Gene expression profiles in pancreatic intraepithelial neoplasia reflect the effects of Hedgehog signaling on pancreatic ductal epithelial cells. Cancer Res. 65 (2005), 1619–26.

33. Ganguly K, et al. Secretory Mucin 5AC Promotes Neoplastic Progression by Augmenting KLF4-Mediated Pancreatic Cancer Cell Stemness. Cancer Res. 1 (81) (2021), 91–102.

34. Cham CM, et al. Glucose deprivation inhibits multiple key gene expression events and effector functions in CD8+ T cells. Eur J Immunol 38 (2008), 2438–2450.

35. Chang CH, et al. Posttranscriptional control of T cell effector function by aerobic glycolysis. Cell 153 (2013), 1239–1251.

36. Michalek RD, et al. Cutting edge: distinct glycolytic and lipid oxidative metabolic programs are essential for effector and regulatory CD4+ T cell subsets. J Immunol 186 (2011), 3299–3303.

37. Connor AA, et al. Pancreatic cancer evolution and heterogeneity: integrating omics and clinical data. Nat Rev Cancer 22 (2022), 131–142.

38. Erickson A, et al. Spatially resolved clonal copy number alterations in benign and malignant tissue. Nature. 608 (7922) (2022), 360–367

39. Liffers ST, et al. Molecular heterogeneity and commonalities in pancreatic cancer precursors with gastric and intestinal phenotype. Gut. 72(3) (2023), 522–534

40. Marusyk A, et al. Intra-tumour heterogeneity: a looking glass for cancer? Nat Rev Cancer 12 (2012), 323–334.

41. Neesse A, et al. Stromal biology and therapy in pancreatic cancer. Gut 60 (2011), 861–868.

42. Waghray M, et al. Deciphering the role of stroma in pancreatic cancer. Curr Opin Gastroenterol 29 (2013), 537–543.

43. Elyada E, et al. Cross-Species Single-Cell Analysis of Pancreatic Ductal Adenocarcinoma Reveals Antigen-Presenting Cancer-Associated Fibroblasts. Cancer Discov 9 (2019), 1102–1123.

44. Ohlund D, et al. Distinct populations of inflammatory fibroblasts and myofibroblasts in pancreatic cancer. J Exp Med 214 (2017), 579–596.

45. Ho WJ, et al. The tumour microenvironment in pancreatic cancer - clinical challenges and opportunities. Nat Rev Clin Oncol 17 (2020), 527–540.

46. Provenzano PP, et al. Enzymatic targeting of the stroma ablates physical barriers to treatment of pancreatic ductal adenocarcinoma. Cancer Cell 21 (2012), 418–429.

47. Clark CE, et al. Dynamics of the immune reaction to pancreatic cancer from inception to invasion. Cancer Res 67 (2007), 9518–9527.

48. Bernard V, et al. Single-Cell Transcriptomics of Pancreatic Cancer Precursors Demonstrates Epithelial and Microenvironmental Heterogeneity as an Early Event in Neoplastic Progression. Clin Cancer Res 25 (2019), 2194–2205.

49. Hiraoka N, et al. CXCL17 and ICAM2 are associated with a potential anti-tumor immune response in early intraepithelial stages of human pancreatic carcinogenesis. Gastroenterology 140 (2011), 310–321.

50. Yao W, et al. Recent insights into the biology of pancreatic cancer. EBioMedicine 53 (2020), 102655.

51. Sun L, et al. IGFBP2 promotes tumor progression by inducing alternative polarization of macrophages in pancreatic ductal adenocarcinoma through the STAT3 pathway. Cancer Lett 500 (2021), 132–146

52. J. Chiou, et al. Follistatin-like protein 1 inhibits lung cancer metastasis by preventing proteolytic activation of osteopontin, Cancer Res. 79 (24) (2019), 6113–6125.

53. Zhao, et al. The role of osteopontin in the progression of solid organ tumour, Cell Death Dis. 9 (3) (2018), 356.

54. Chen, et al. Single cell RNA-seq identifies immune-related prognostic model and key signature-SPP1 in pancreatic ductal adenocarcinoma, Genes 13 (10) (2022).

